# Inhibition Controls Receptive Field Size, Sensitivity, and Response Polarity of Direction Selective Ganglion Cells Near the Threshold of Vision

**DOI:** 10.1101/683961

**Authors:** Xiaoyang Yao, Greg D. Field

## Abstract

Information about motion is encoded by direction-selective retinal ganglion cells (DSGCs). These cells reliably transmit this information across a broad range of light levels, spanning moonlight to sunlight. Previous work indicates that adaptation to low light levels causes heterogeneous changes to the direction tuning of ON-OFF (oo)DSGCs and suggests that superior-preferring ON-OFF DSGCs (s-DSGCs) are biased toward detecting stimuli rather than precisely signaling direction. Using a large-scale multi-electrode array, we measured the absolute sensitivity of ooDSGCs and found that s-DSGCs are ten-fold more sensitive to dim flashes of light than other ooDSGCs. We measured their receptive field sizes and found that s-DSGCs also have larger receptive fields than other ooDSGCs, however, the size difference does not fully explain the sensitivity difference. Using a conditional knockout of gap junctions and pharmacological manipulations, we demonstrate that GABA-mediated inhibition contributes to the difference in absolute sensitivity and receptive field size at low light levels, while the connexin36-mediated gap junction coupling plays a minor role. We further show that GABA-mediated inhibition masks the OFF response of ooDSGCs under scotopic conditions, restricting their responses to increases in light. These results reveal that GABAergic inhibition controls and differentially modulates the responses of ooDSGCs under scotopic conditions.

**Significance Statement:** Light adaptation and parallel processing are two major functions of retina. Here we show that parallel processing is differentially regulated between photopic and scotopic conditions across DSGCs. This differential adaptation alters the absolute sensitivity and RF size of s-DSGCs relative to other ooDSGC types. These results point to novel mechanisms and possibly new circuit elements that shape retinal processing of motion under rod-mediated light levels.

## Introduction

Detecting motion in the environment while accurately estimating its direction and speed are major functions performed by both vertebrate and invertebrate visual systems (Borst et al., 2010; Mauss et al., 2017; Wei, 2018). In mammals, motion and its direction are computed and recomputed at several stages of visual processing (Barlow et al., 1964; Born and Bradley, 2005; Hillier et al., 2017), illustrating the importance of these signals for vision. The first cells that explicitly report motion direction are in the retina. These direction-selective ganglion cells (DSGCs) exhibit responses that are tuned to one of four ‘cardinal’ directions: superior, inferior, anterior or posterior. A major class of DSGCs, the so-called ON-OFF (oo)DSGCs, exhibit broad speed tuning and send axons to multiple brain areas including the superior colliculus and lateral geniculate nucleus (Dhande and Huberman, 2014; Gauvain and Murphy, 2015; Huberman et al., 2009). Thus, these neurons likely contribute to a range of behaviors that rely on rapidly processing motion in the environment (Sabbah et al., 2017; Sun et al., 2015).

To reliably signal motion between a moonless night and a sunny day, animals must accommodate a change in the mean photon flux that spans twelve orders of magnitude (Rodieck, 1998). This presents a challenge to the retina to maintain high sensitivity while avoiding saturation. Previous studies show that ooDSGCs exhibit two major changes in their responses between scotopic (rod-mediated) and photopic (cone-mediated) lighting conditions. First, under photopic conditions, ooDSGCs exhibit roughly equal responses to objects brighter or dimmer than the background. However, under scotopic conditions, they lose their ‘OFF’ response and respond primarily to increments of light (Pearson and Kerschensteiner, 2015). The second identified form of adaptation is a broadening of direction tuning among superior-preferring (s-)DSGCs at low light levels: the other three ooDSGC types preserve their tuning widths across light levels (Yao et al., 2018). This change in the relative tuning has been explained as an adaptive strategy that sacrifices some information about the direction of motion in order to improve the ability of the population to signal the presence of motion (Yao et al., 2018). This explanation suggests that s-DSGCs may be more sensitive to just-detectable stimuli as they play a dominant role in signaling the presence of a stimulus, while the other types play a dominant role in signaling its direction of motion.

To test this idea, we presented just-detectable, full-field flashes of light to the ex vivo mouse retina while recording the responses of ooDSGCs using a large-scale multielectrode array (MEA). We found that s-DSGCs were ~10-fold more sensitive to flashes of light near absolute visual threshold than the other three ooDSGC types. Furthermore, s-DSGC sensitivity was within a factor of three of ON-alpha cell sensitivity, which is likely one of the most sensitive RGC types in the mouse retina ((Takeshita et al., 2017), but see (Pan et al., 2016)). One possibility for why s-DSGCs are more sensitive than other ooDSGCs is that they integrate signals over a larger area. We tested this possibility by measuring spatial receptive field (RF) sizes and their dependence on gap-junction coupling (s-DSGCs are the only DSGC type that is homotypically coupled). Neither RF size nor gap junction coupling could account for the sensitivity difference. We then tested the role of GABAA-mediated inhibition in tuning response sensitivity, because previous work suggested (1) differences between the inhibition impinging on s-DSGCs relative to other ooDSGC types under scotopic conditions (Yao et al., 2018), and (2) that inhibition controls the absolute sensitivity of many RGCs (Pan et al., 2016). We found that application of gabazine made the sensitivity difference across DSGC types much smaller by greatly increasing the flash sensitivity of inferior, posterior, and anterior preferring ooDSGC. Furthermore, application of gabazine revealed that inhibition controls the RF sizes of ooDSGCs and suppresses their OFF responses under scotopic conditions. These results reveal several novel facets to GABA-mediated inhibition shaping ooDSGC response polarity, RF size, and absolute sensitivity near visual threshold. These results also suggest a difference in the functional circuitry providing input to s-DSGCs vs other ooDSGC types under scotopic conditions.

## Materials and Methods

### Animals

All animals were healthy adults between the age of 2 months and 1 year, and housed in 12hr light-dark cycles, in groups up to 5 animals per cage with *ad lib* access to food and water. Both male and female mice were used in the study. All procedures to maintain and use mice were in accordance with the Duke University Institutional Animal Care and Use Committee. C57BL/6J (RRID: IMSR_JAX:000664) mice were acquired from The Jackson Laboratory. FACx mice were acquired from Dr. Gautam Awatramani at University of Victoria and have been described previously (Yao et al., 2018).

### Recording procedures

Animals were dark-adapted overnight and euthanized by decapitation in the dark using infrared converters. Following euthanasia, the eyes were removed and a piece of dorsal peripheral retina was isolated and mounted on an array of extracellular microelectrodes (MEA) with ganglion cell side down (Yao et al., 2018). To facilitate the delivery of visual stimuli that drove photoreceptor responses, dorsal retina was used to measure responses from areas where cone photoreceptors exhibit the highest expression of middle-wavelength sensitive opsin (Applebury et al., 2000; Wang et al., 2011). The MEA consists of 519 electrodes with 30 µm spacing, covering a hexagonal region 450 µm across (Field et al., 2010; Yao et al., 2018). While recording, the retina was perfused with Ames’ solution (30-32°C) bubbled with 95% O_2_ and 5% CO_2_, pH 7.4.

### Spike sorting and EI analysis

Recordings were analyzed offline to identify and sort the spikes of different cells into different clusters (Dabrowski et al., 2004; Field et al., 2007; Yu et al., 2017). Briefly, individual spikes were identified by voltage thresholding on every electrode. The spike waveforms were extracted and clustered using a mixture of Gaussian model in a five-dimensional space determined by principal components analysis. Clusters with more than 10% contamination in refractory period or less than 100 spikes were excluded. Neurons sharing >25% of their spike times in common were considered duplicates and the copy with the fewer number of spikes was excluded from further analysis.

An electrical image (EI) was calculated by averaging voltage on each electrode 1 ms preceding and 3 ms following a spike, reflecting the cell’s position, extent of dendritic arbor and axon trajectory relative to the electrode array. The orientation of the dorsal-ventral axis in each recording was determined by the direction axons traveled from each RGC in the EIs (Yao et al., 2018). EIs were also used to track individually recorded neurons across light levels and stimulus conditions (Field et al., 2009; Yao et al., 2018).

### Light stimulation

The retina was stimulated with a gamma-corrected OLED microdisplay refreshing at 60.35 Hz (SVGA+XL Rev3, eMagin. Santa Clara, CA). The stimuli were created in MATLAB (Mathworks, Natick, MA) and projected through the microscope objective (Nikon, CFI Super Fluor 4x) and focused onto the photoreceptor outer segments. The emission spectrum of each display primary was measured with a PR-701 spectra-radiometer (PhotoResearch) after passing through the optical elements between the display and the retina. Photon flux was measured as described before to calibrate the display (Yao et al., 2018). Three classes of stimuli were used:

#### Flashing squares

Flashing squares were used to measure spatial RFs of DSGCs. A single positive-contrast square was turned on for 1 s and turned off for 1 s at one location and presented pseudo-randomly at different (non-overlapping) locations over the whole stimulation field. The size of a stimulus square was between 60μm X 60μm and 160μm X 160μm. The square had 900% Michelson contrast to elicit robust responses. To accurately estimate RF sizes, we used flashes spanning a larger area to measure s-DSGC RFs and smaller area flashes for the other ooDSGC types (Fig 2D-E); this could bias the RF size estimates. As a control, we compared RF sizes across cell types with the same flash sizes and found that estimated RF sizes did not vary significantly between flashed squares that were 60 μm to 120 μm on an edge.

#### Full-field flashes

The second stimulus type used were full field flashes to measure the absolute sensitivity of DSGCs. Flashes were presented in complete darkness. The flash duration varied between 2-8 ms, which together with neutral density filters modulated the mean number of delivered photons. A flash of certain strength was delivered every three seconds for 60 repetitions. Flashes with different strengths were presented in order of lowest to highest strength. Responses to these full-field flashes were used in a 2AFC discrimination task and ideal observer analysis (see below) to distinguish ‘flash’ and ‘no flash’ epochs. The ‘no flash’ epochs were obtained by collecting three minutes of spontaneous activity at the beginning of each experiment and dividing these into 60, three-second trials.

#### Drifting gratings

Drifting gratings were used to distinguish DSGCs from non DSGCs using classification methods described previous (Yao et al., 2018). Gratings moved in 8 directions with speeds ranging from 24-2400 μm/s, at a spatial period of 960 μm (49.2° visual angle) and at 50% Michelson contrast. Drifting gratings of different directions and speeds were presented in a pseudo-random order. Each grating was presented for 8 seconds with 2 seconds of gray screen between presentations. Each grating was presented 3-4 times, and responses were averaged over presentations. ooDSGCs were distinguished from ON DSGCs based on their speed tuning (Yao et al., 2018).

### Two-alternative forced-choice analysis

The absolute sensitivity of each DSGC was quantified as the fraction of correctly discriminated trials in which a flash was presented (‘flash’ trials) from trials during which no stimulus was presented (‘no flash’ trials). The discrimination was performed using a 2AFC analysis (Chichilnisky and Rieke, 2005; Dhingra and Smith, 2004; Field et al., 2019). The response in each trial was summarized by a vector, which specified the spike count as a function of time (in 20 ms bins). Using a subset of the data at each flash strength, a discriminant was calculated by taking the difference between the mean ‘flash’ trials and the mean of the ‘no flash’ trials. A different discriminant was calculated at each flash strength to optimize the discriminant for response latencies and dynamics that changed across flash strengths. Using a different subset of the data, the dot product between the discriminant was calculated for both a randomly selected ‘flash’ trial and a randomly selected ‘no flash’ trial. The dot product with the larger value was classified as the ‘flash’ trial. This procedure was iterated across all ‘flash’ and ‘no flash’ trials without replacement. Performance was calculated as the fraction of correctly classified trials. This ‘difference of means’ discrimination procedure assumes independent and additive noise with no covariance (Duda et al., 2000) and has been shown previously to work well in flash detection tasks applied to salamander RGCs (Chichilnisky and Rieke, 2005). Assuming, independent and additive Gaussian noise, the relation between discrimination performance (probability correct) and signal-to-noise ratio (SNR) is:

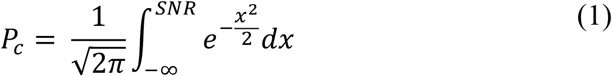

P_c_ = 0.84 was set to be the detection threshold when SNR = 1. Cells with sensitivity curves never exceed 0.84 were excluded from the analysis.

The discrimination performance curves were fitted to the Naka-Rusthton equation: 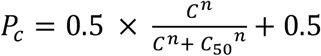, where C indicates the flash intensity, C_50_ the semi-saturation constant and n is a coefficient proportional to the slope of the sensitivity curve at C_50_.

### Rod photoreceptor simulation and pooling model

A model of rod responses and was used to test how differences in RF size across DSGCs impacts their predicted thresholds to just detectable flashes in a 2AFC task. The simulation and pooling of rod signals was examined by using a recently published model of rod responses (Field et al., 2019). The model includes specific factors to capture the three dominant noise sources in the rod photocurrent: continuous fluctuations in the dark current (a.k.a. continuous noise) (Baylor et al., 1980; Baylor et al., 1984), discrete noise events in the dark-current produced by the thermal activations of rhodopsin, and variability in the amplitude and kinetics of the single photon response (Baylor et al., 1979; Field and Rieke, 2002a; Rieke and Baylor, 1998). A rod response, *r(t)* was generated from the following equation:

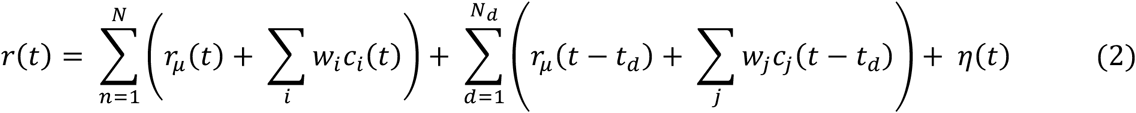

Continuous dark noise, *η*(*t*), was generated by sampling from a Gaussian distribution and filtering in time to match the power spectrum of measured continuous noise (Field et al., 2019). *N* photon responses in an individual rod were generated by the first term in Eq. 2 by sampling from a Poisson distribution with a mean given by the flash strength on a given trial. The mean single photon response is given by *r*_*μ*_(*t*), and the covariance of the single photon response is captured by summing over a weighted set, *w*_*i*_, of eigenvectors, *c*_*i*_(*t*), derived from the covariance matrix of the single photon response (Field et al., 2019). Finally, discrete noise events caused by the thermal activation of rhodopsin were captured by the second term of Eq. 2: *N*_*d*_ thermal isomerizations where generated using the same formulation as for the single photon response, but *N*_*d*_ was determined by sampling from a Poisson distribution with a mean given by the thermal isomerization rate and each isomerization event occurred at random and independent times given by *t*_*d*_.

In the original formulation of this model (Field et al., 2019), the noise parameters were derived from measurements of the photocurrent from primate rods. We adjusted these parameters to account for the greater amounts of continuous noise and variability in the single-photon response present in mouse rods (both relative to the amplitude of the mean single-photon response). Specifically, the continuous noise was increased by 22% and single-photon response variability was increased by 37.5% (Field and Rieke, 2002a, b). The kinetics of the single photon response and the shape of the power spectrum of the continuous noise are both similar between mouse and primate rods, so no adjustments were made to these quantities. The thermal rate in mammalian and mouse rods is somewhat uncertain, with the literature allowing for a wide range of values between 0.001 and 0.015 Rh*/rod/s, so we chose an intermediate value of 0.005 Rh*/rod/s (Burns et al., 2002; Field et al., 2019; Fu et al., 2008; Yue et al., 2017). Importantly, this model for simulating rod responses reproduces rod detection and temporal sensitivities in a 2AFC task (Field et al., 2019), supporting the application of this model for analyzing task performance of pools of rods.

For linear pooling, the optimal linear discriminant was generated from a set of 5000 simulated single photon responses (including dark noise), and 5000 simulated rod responses that contained only dark noise: a linear discriminant, *D*, was computed as the difference between the means of these two ensembles: *μ*_*flash*_ and *μ*_*null*_.

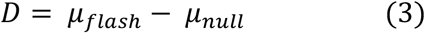

We then simulated individual rod responses for a pool of *P* rods by iterating over Eq 2 for given a flash strength, *f*. The dot product between individual rod responses and the discriminant was computed and summed over all responses to yield an integer value, *R*_*f*_.

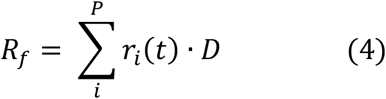

We also simulated individual rod responses for the same size pool of *P* rods for a flash strength of 0 (the ‘no flash’ or ‘null’ case). Again, the dot product between individual responses and the discriminant was computed and summed over all responses as in Eq 4 to yield an integer value, *R*_0_. If *R*_*f*_ > *R*_0_, the trial was scored as correct, if *R*_*f*_ < *R*_0_, the trial was scored as incorrect and if *R*_*f*_ = *R*_0_, the trial was randomly scored as correct with 50% probability. This procedure was iterated over 500 trials to calculate the probability correct in the 2AFC task for flash strength *f*.

For nonlinear pooling, the discriminant and simulated rod responses were generated in the same manner as described for linear pooling. The only difference was that after computing the dot-product of a simulated rod response with the discriminant, the result was weighted by the likelihood ratio between the probability that the response arose from the distribution of single photon responses versus the distribution of continuous noise. These probabilities were estimated by generating two separate response ensembles: a training set of 5000 simulated single photon responses and another set of 5000 responses containing only continuous noise. The dot product of each of these response ensembles with the linear discriminant was computed and were nearly Gaussian. Thus, their means and standard deviation were used to summarize the distributions: *μ*_*A*_, *μ*_*B*_, *σ*_*A*_, and *σ*_*B*_, where *A* and *B* denote the distributions of flash and no flash responses, respectively, after computing their dot products with the discriminant. The optimal (Bayesian) nonlinear weighting was thus calculated as

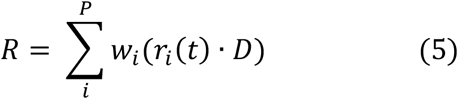

where

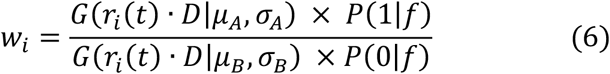

where *G*(*X*|*μ*,*σ*) is the probability of sampling X from a Gaussian distribution with mean, *μ*, and standard deviation *σ*; *P*(*Y*|*λ*) is the probability of sampling *Y* from a Poisson distribution with mean *λ*; and *f* in Eq 6 is the flash strength, which acts like a prior in the Bayesian sense.

Responses were simulated for flash strength *f* and compared to responses of flash strength 0. As with linear pooling, if *R*_*f*_ > *R*_0_, the trial was scored as correct, if *R*_*f*_ < *R*_0_, the trial was scored as incorrect and if *R*_*f*_ = *R*_0_, the trial was randomly scored as correct with 50% probability.

The simulated rod pool sizes were estimated from the RF sizes of the ooDSGCs as follows: Given a Gaussian RF size with a standard deviation of 0.012 mm^2^ for ooDSGCs (Fig 2, excluding s-DSGCs), the equivalent pool size for uniformly weighted rod signals is twice as large (0.024 mm^2^)(Hemila et al., 1998). The number of rods in this RF is 12,000, given a rod density of 500,000 rods/mm^−2^ in mouse (Jeon et al., 1998; Volland et al., 2015). s-DSGCs were taken to be eight times larger: 96,000 rods.

### Statistical analysis

Unless otherwise noted, population averages are expressed as mean ± SEM. The number of cells and retinas are indicated in the figure legends; n represents the number of cells used unless indicated otherwise. Each retina came from a different mouse. Student’s t test was used to compare values under different conditions, and the differences were considered significant when p ≤ 0.05.

## Results

### Superior DSGCs are 10-fold more sensitive to detecting dim flashes

In a recent study, we showed that s-DSGCs exhibit broader direction tuning and higher response gain at low light levels (Yao et al., 2018). This suggests that s-DSGCs may be more sensitive to just-detectable stimuli than the other three ooDSGC types under scotopic conditions.

To test this, we compared the sensitivity of different types of RGCs under dark-adapted conditions to dim flashes of light. Brief (2-8 ms), full-field flashes were delivered to the mouse retina, *ex vivo*, while recording the spikes of RGCs over an MEA (Fig 1A). Flashes ranged from strengths that rarely elicited spikes (0.001 Rh*/rod) to strengths that yielded robust spiking (1 Rh*/rod). Spike rasters of individual RGCs indicated that s-DSGC responded robustly to weaker flash strengths than the other ooDSGC types (Fig 1A).

**Figure 1:**
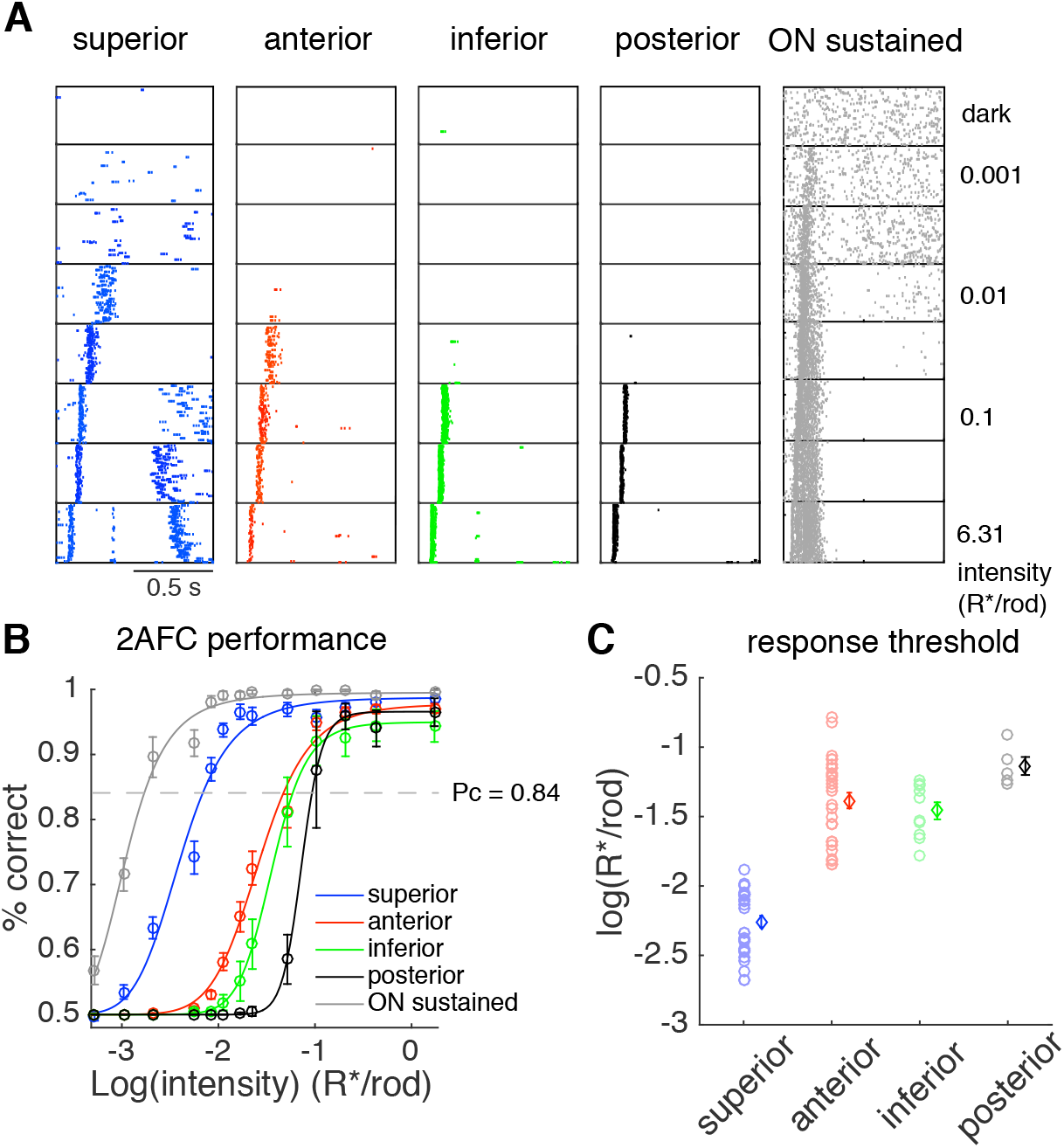
Superior DSGCs have a lower absolute detection threshold than other ooDSGCs. **A.** Rasters show spike responses of example cells to brief, full-field flashes (2-8 ms) in darkness. Top row shows spontaneous activity. Second to bottom rows shows responses to flashes with intensity increasing from 0.001 to 6.31 R*/rod. Flashes were presented at the left edge of each panel. For each intensity, flashes were repeatedly delivered every 3 seconds, 60 times. **B.** Discrimination performance in two-alternative forced-choice (2AFC) task, as a function of flash intensity. The dash line indicates 83% correct in the task (detection threshold). Data were fitted by the Naka-Rushton equation (See Method). Data were collected from one retina (Superior: n = 16, Anterior: n = 17, Inferior n = 8, Posterior: n = 5, ON Sustained: n = 11). **C.** Detection threshold of different DSGC types. Each circle shows detection threshold of an individual cell. Each diamond represents the mean ± SEM threshold of a cell type. Data from 2 retinas (Superior: n = 31, Anterior: n = 29, Inferior: n = 10, Posterior: n = 5).

To quantify and compare the sensitivity to dim flashes across RGCs, a two-alternative forced-choice (2AFC) task was performed on spike trains to classify responses as arising from ‘flash’ or ‘no flash’ trials (Chichilnisky and Rieke, 2005). The response classification was based on a linear discriminant learned from training responses and applied to test data (see Methods). This analysis resulted in a neurometric sensitivity curve (Fig 1B), quantifying the accuracy with which a response predicts the occurrence of a flash over a range of flash intensities. We used the flash intensity that elicited 84% correct in the 2AFC task as detection threshold; this corresponds to a signal-to-noise ratio (SNR) = 1 (Chichilnisky and Rieke, 2005)(See Methods).

We found that the detection thresholds of s-DSGCs were ~10-fold lower than the other three ooDSGC (Fig 1C, p < 0.001), indicating ~10-fold higher sensitivity. Furthermore, the thresholds of the s-DSGCs approached the lowest thresholds observed among any of the recorded RGCs over the MEA (Fig 1A, gray). These were RGCs with sustained ON responses and large spatial RFs (data not shown): these RGCs are likely ON sustained alpha-cells because they exhibited large spikes over the MEA, presumably produced by large somata (Ravi et al., 2018; Takeshita et al., 2017; Yu et al., 2017). These observations indicated that s-DSGCs are more sensitive to dim flashes of light than the other ooDSGC types and approach (within ~0.5 log unit) the sensitivity of the most sensitive RGCs in the dark-adapted retina. Below we investigate the mechanisms that produce the higher sensitivity of s-DSGCs than other ooDSGC types under dark-adapted conditions.

### Superior ooDSGCs exhibit larger spatial RFs than other ooDSGC types

One possible explanation for the lower thresholds exhibited by s-DSGCs than the other ooDSGC types is that they have larger RFs, thereby having a larger area in which to collect more photons. To measure the RF size of ooDSGCs, we presented spatially restricted flashes. Each flash was presented at a psuedorandom and non-overlapping location on the retina; the flash was turned ‘on’ for one second with a one-second inter-stimulus interval. The one-second presentation allowed separating ON from OFF responses (Fig 2A, blue and red, respectively) and thus separating the spatial RFs into ON and OFF subfields (Fig 2B-C). The area of the ON and OFF subfields were then estimated by fitting each with a two-dimensional Gaussian function and computing the area encompassed by the one-sigma contour.

**Figure 2.**
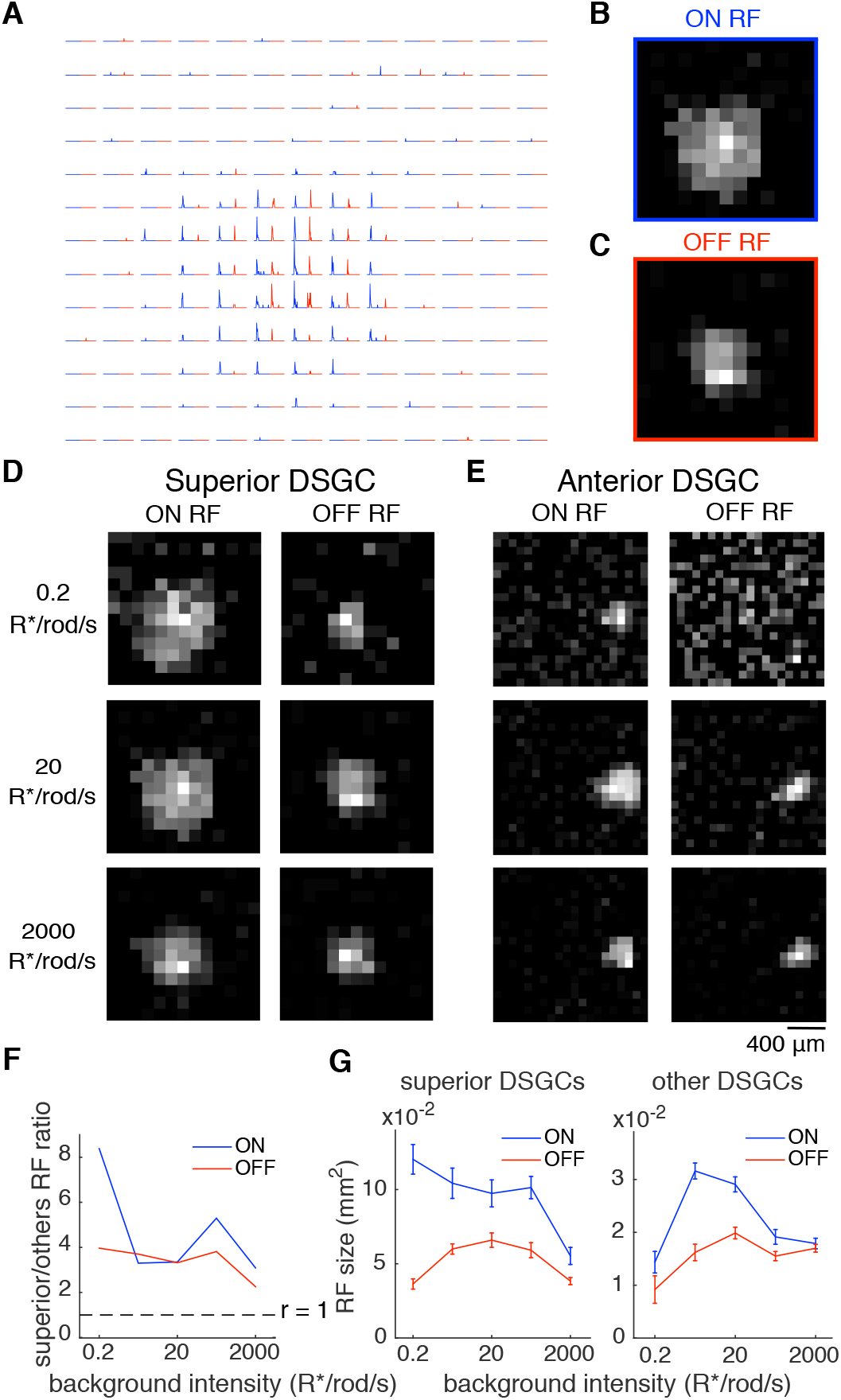
Receptive field sizes of DSGCs depend on mean light level. **A.** Spiking response of an example DSGC to onset (blue) and offset (red) of flashed squares at each of 169 locations on a 13 × 13 grid. Stimuli were turned on at the beginning of each trace for 1 second, and then turned off for one second. Data were averaged across 5 trials for each location. **B-C.** ON (B) and OFF (C) subfields for the cell shown in A. Heat maps calculated from the average spike number at each location normalized to the peak response across the whole field. **D.** RF of an example superior DSGC across 3 light levels. Left: ON subfield. Right: OFF subfield. The background light intensity from top to bottom are: 0.2, 20, 2,000 R*/rod/s. The size of the flashing square is 120 × 120 μm. **E.** RF of an example anterior DSGC across 3 light levels. The size of the flashed square is 80 × 80 μm. **F.** Ratio between superior and other DSGC RF sizes as a function of background light intensity. The dash line indicates equal size of RFs of the two groups. **G.** Average RF size of DSGCs is plotted as a function of background light intensity. RF sizes were estimated from Gaussian models fitted to heat maps shown in B-E. Left: superior DSGCs (n = 23). Right: combined other three directions (n = 30, anterior: n = 19, inferior: n = 6, posterior: n = 5).

The RF sizes of s-DSGCs were larger than the other three types over a range of light levels spanning scotopic and photopic conditions (Fig 2D-G; p < 0.001). The difference was largest at the lowest light level we tested (ON subfield: ratio = 8.4, OFF subfield: ratio =4.0 at 0.2 Rh*/rod/s) and smaller at higher light levels (ON subfield: ratio ≈ 3.1, OFF subfield: ratio ≈ 2.3 at 2000 Rh*/rod/s). Among the other three types of DSGCs, there was no statistical difference in their mean RF sizes at most light levels (P > 0.05, data not shown). These data illustrate that s-DSGCs exhibit larger RFs than the other ooDSGCs and that this difference is largest under low scotopic conditions.

### Difference in RF size cannot account for sensitivity difference

Given the ~8-fold larger ON subfields of s-DSGCS than other ooDSGCs at 0.2 Rh*/rod/s (Fig 2F), we examined the extent to which this size difference could account for the sensitivity difference in the 2AFC task (Fig 1). Under a simple model that assumes linear pooling of rod responses with independent and additive Gaussian noise, an 8-fold increase in the number of rods will produce an 8-fold increase in signal and √8-fold increase in noise. This results in a √8-fold increase in the signal-to-noise ratio (SNR). Thus, this simple linear pooling model predicts ~3-fold increase in the 2AFC performance, significantly less than the ~10-fold difference that we observed (Fig 1).

This simple analysis has two limitations. First, the dominant source of noise in rods, the thermal activation of rhodopsin, is not a noise source with a zero-mean amplitude distribution. This is because these noise events masquerade as photon responses (Yau et al., 1979). Second, several lines of evidence indicate that the retina is *nonlinearly* pooling rod responses (Berntson et al., 2004; Field and Rieke, 2002b). Thus, we also examined a more complex model in which all sources of rod noise were accurately modeled and rod signals were optimally and nonlinearly pooled (see Methods) (Field et al., 2019). For s-DSGCs and other ooDSGCs, we simulated rod responses in pools of 96,000 and 12,000 rods, respectively (see Methods). We then performed the same 2AFC discrimination task on these simulated responses as we performed on DSGC responses. While the overall performance of nonlinear pooling exceeded a linear pooling model, we observed only a 70% increase in performance between pools of 96,000 and 12,000 rods. This is because the thermal activation of rhodopsin is an asymmetric noise source that does not average to zero as more rods are added to the pool. This results in diminishing returns in sensitivity with larger RF sizes.

To summarize, given just an ~8-fold difference in RF size, both a linear and an optimal nonlinear pooling models predict < 3-fold difference in sensitivity -- much less than the ~10-fold observed sensitivity difference. We conclude from these analyses that the difference in RF size cannot explain the difference in absolute sensitivity.

### Receptive field sizes of ooDSGCs depend on light level and direction preference

Before testing potential mechanisms that could account for the lower detection thresholds of s-DSGCs, we examined the dependence of ooDSGC RF size on light level. Previous studies have indicated a relatively monotonic relationship between the RF center sizes of non-DS RGCs, with larger RFs under scotopic than photopic conditions (Barlow et al., 1957; Muller and Dacheux, 1997; Troy et al., 1999). Do ooDSGC RF sizes change monotonically with light level?

For s-DSGCs, ON subfields were largest under scotopic conditions, and decreased in a relatively monotonic fashion at higher light levels (Fig 2G, left). However, the OFF subfields of s-DSGCs were largest under mesopic conditions (Fig 2G, left). Among the other ooDSGC types, ON subfields were largest under mesopic conditions and smaller under low scotopic and photopic conditions, while OFF subfields did not change much from mesopic to photopic conditions, but were smaller under scotopic conditions (Fig 2G, right). Finally, across all ooDSGC types, ON subfields were larger than OFF subfields at low light levels but were nearly equal under photopic conditions (Fig 2G). Cumulatively, these results suggest that the underlying circuit mechanisms mediating adaptation are non-uniform across ON and OFF subfields and also differ between s-DSGC and other ooDSGC types. We note that RF sizes likely depend on stimulus strength, with larger contrast changes potentially eliciting responses that rise above threshold near the edge of the RF. The small spots of light we used to probe the RF size were 900% contrast above the background illumination, which is likely to drive responses even near the edge of the RF.

Furthermore, natural scenes rarely contain contrasts above this value (Mante et al., 2005). Thus, given these high contrast probe stimuli, we do not think we are underestimating the RF size under contrasts experienced in typical viewing conditions.

### Absolute sensitivity and receptive field size are modestly affected by gap junctions

Given s-DSGCs exhibit lower thresholds (Fig 1) and larger RFs under scotopic conditions than other ooDSGC types (Fig 2), we sought to identify the underlying mechanisms that cause these differences. One of the major difference between s-DSGCs and other ooDSGCs is that s-DSGCs are homotypically coupled with their neighbors via connexin36-mediated gap junctions: other ooDSGC are not homotypically coupled (Trenholm et al., 2013; Trenholm et al., 2014; Vaney, 1994). Previous studies have shown that these gap junctions contribute to several properties of s-DSGCs, including broader tuning curves at low light levels (Yao et al., 2018), and priming responses for moving stimuli (Trenholm et al., 2013). To test for a contribution of gap-junctions to absolute sensitivity and RF size, we used FACx mice: a conditional knockout line in which the expression of the connexin36 gene is selectively disrupted in s-DSGCs (Yao et al., 2018). If homotypic coupling plays a significant role in the higher sensitivity of s-DSGCs, then the sensitivity should be reduced (threshold increased) among s-DSGCs in FACx mice. These mice were used instead of the constitutive Cx36 knockout to avoid disrupting gap junction coupling in the circuits upstream to s-DSGCs, including homotypic coupling among AII amacrine cells and heterotypic coupling between AII amacrine and cone bipolar cells (Bloomfield and Dacheux, 2001).

First, we measured the absolute sensitivity of ooDSGCs in dark-adapted FACx retinas (Fig 3A, see Methods) and compared it to the sensitivity in WT (C57/BL6-J) retinas (Fig 3B). Similar to WT, s-DSGCs in the FACx retinas exhibit detection thresholds ~10-fold lower than the other three ooDSGC types (Fig 3A). Furthermore, there was no significant difference in the absolute thresholds between WT and FACx retinas for either s-DSGCs (p = 0.61) or other ooDSGCs (p = 0.32; Fig 3B). Thus, Cx36 gap junctions did not contribute significantly to the higher absolute threshold of s-DSGCs.

**Figure 3.**
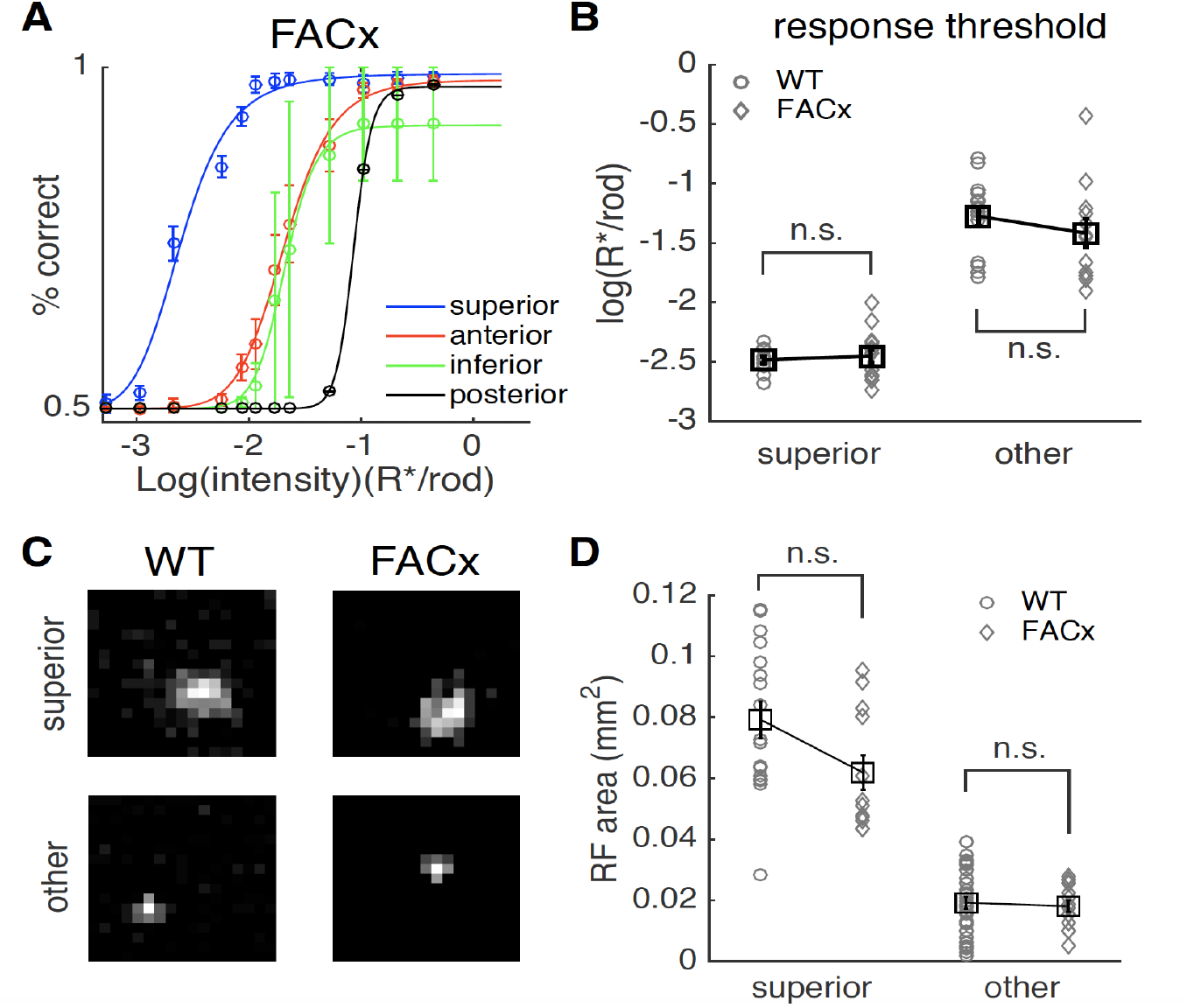
Effect of gap junction coupling on absolute sensitivity and RF size. **A.** Discrimination performance of DSGCs in FACx mice (one retina, Superior: n = 14, Anterior: n = 9, Inferior: n = 2, Posterior: n = 1). Data were fitted by the Naka-Rushton equation (See Method). **B.** Detection threshold of different DSGC types in C57 and FACx mice (C57: one retina, s-DSGCs n = 14, other DSGCs n = 15). Discrimination performance curve of individual cells were fitted by the Naka-Rushton equation. Each gray circle or diamond represents detection threshold of an individual cell. Each square represents the mean +- SEM threshold of a cell type. **C.** Example RFs of s-DSGCs and other DSGCs from C57 and FACx mice. The RF sizes were estimated from summing the ON- and OFF- responses. Stimulus square size: 120μm X 120μm, background light intensity: 0.2 R*/rod/s. **D.** Comparison of RF size between s-DSGCs and other DSGCs in C57 and FACx mice (C57 s-DSGCs: n = 17, FACx s-DSGCs: n = 12, C57 other DSGCs n = 37, FACx other DSGCs n = 15). The RF sizes were estimated from summing the ON- and OFF- responses.

Next, we compared the RF size of DSGCs between FACx and WT retinas under scotopic conditions (1 Rh*/rod/s). Gap junction coupling could potentially increase the RF size of RGCs by pooling photoreceptor signals from a larger area (Shelley et al., 2006). However, we found that the RF sizes of s-DSGCs modestly changed in the FACx retinas relative to WT (Fig 3C-D: p = 0.052) while the RFs sizes of other ooDSGCs did not change (Fig 3C-D: p = 0.5). We also compared ON- and OFF- subfield sizes between FACx and WT retinas for all ooDSGC types and found no significant difference (data not shown). These results indicate that homotypic gap junction coupling does not produce the difference in absolute sensitivity or RF size between s-DSGCs and other ooDSGCs types. The weak impact of gap junctions on threshold and RF size may be explained by two factors: (1) Gap junction coupling is expected to play a minor role when coupled cells respond to a homogenous full field stimulus, because the voltage differences among coupled cells will be small. (2) The gap junction conductance in s-DSGCs is small and insufficient to elicit spikelets without coinciding with chemical synaptic input (Bloomfield and Xin, 1997; Trenholm et al., 2014; Trong and Rieke, 2008).

### GABA_A_ antagonists reduce sensitivity difference and increase RF size among DSGC types

An alternative mechanism controlling the absolute sensitivity of DSGCs is the effective inhibition that they receive. Previous studies suggest that response thresholds of RGCs fall into distinct groups in mouse, with low, medium, and high thresholds (Volgyi et al., 2004). Furthermore, the threshold differences that separate these groups can be mitigated by pharmacologically blocking GABAergic inhibition (Pan et al., 2016). Thus, we hypothesized that the distinction among response thresholds in different ooDSGC types could be caused by differences in effective GABAergic inhibition. To test this possibility, we used the GABA_A_ receptor antagonist gabazine (SR-95531) to attenuate inhibitory input onto DSGCs and their presynaptic circuits (Pei et al., 2015; Yoshida et al., 2001)(see Methods). Application of gabazine increased the spontaneous firing rate for DSGCs in darkness (Fig 4A), increasing the background noise upon which dim flashes had to be detected. However, our use of a 2AFC task and ideal observer analysis to classify responses as ‘flash’ or ‘no flash’ trials revealed a net increase in the signal-to-noise ratio, because detection thresholds were lower following application of gabazine (Fig 4A-C, p < 0.001). Although there was some residual difference in response threshold between s-DSGCs and other ooDSGC types (p < 0.001), the difference decreased from 9.4-fold, on average, in control to 3.2-fold, on average, in gabazine (Fig 4D). We also applied the GABA_C_ receptor antagonist TPMPA but did not find any further change in the response threshold of DSGCs (data not shown). These results indicate that GABA_A_ receptor mediated inhibition largely controls the absolute sensitivity of different types of DSGCs (see Discussion).

**Figure 4.**
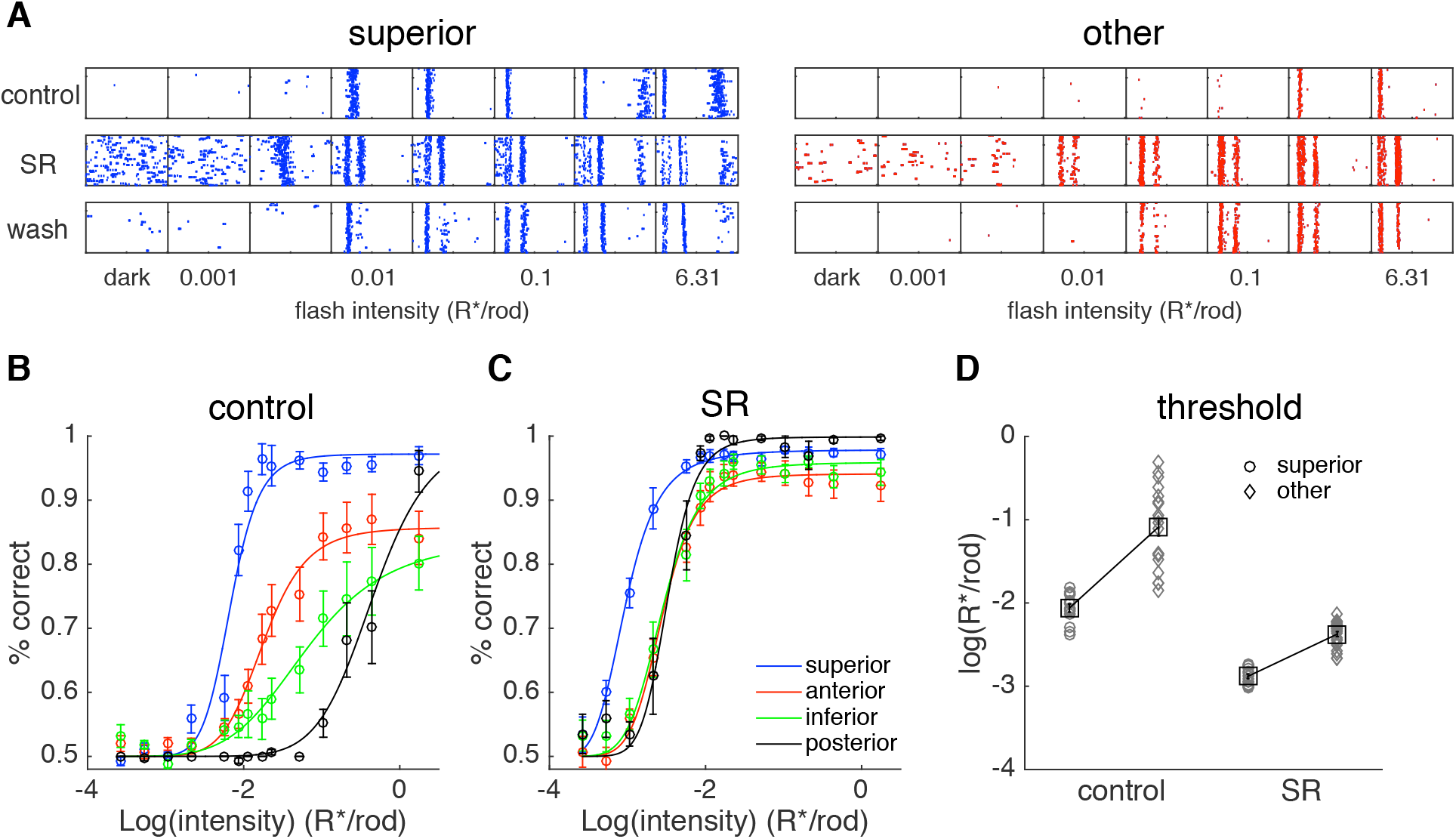
GABA_A_ blocker decreases absolute threshold of DSGCs. **A.** Rasters show spike responses of example superior and other DSGCs to brief dim full field flashes (2-8 ms) in darkness under control, SR (15μM) and wash conditions. The leftmost column of each cell shows spontaneous activity. Second from left to last columns show responses to flashes with intensity increasing from 0.001 to 6.31 R*/rod. Flashes were repeatedly delivered every 3 seconds, 60 repeats. SR was washed in for 5 minutes and washed out for 30 minutes. **B-C.** Discrimination performance of C57 DSGCs in control and SR conditions (one retina, Superior: n = 15, Anterior: n = 15, Inferior: n = 10, Posterior: n = 5). Data were fitted by the Naka-Rushton equation (See Method). **D.** Detection threshold of different DSGC types under control and SR conditions. Discrimination performance curves from individual cells were fitted by the Naka-Rushton equation, and the flash strength producing 83% correct (threshold) performance was extracted from the fit. Each circle or diamond represents detection threshold of an individual cell. Each square represents the mean +- SEM threshold of a cell type.

We next examined the impact of gabazine on the RF sizes of s-DSGCs and other ooDSGC types. We again measured RF sizes of ooDSGCs using small-area, 1-second flashes, but now with gabazine. The RF areas of all ooDSGC types increased following application of gabazine (Fig 5A-B). ON subfields exhibited a 40% and 75% increase in area for s-DSGCs and other ooDSGCs, respectively. Surprisingly, there was a much larger fractional increase in RF area for the OFF subfields than the ON subfields under scotopic conditions (Fig 5A-B). OFF subfield exhibited 10-fold and 30-fold increases in area following application of gabazine for s-DSGCs other ooDSGCs, respectively (Fig 5B, right). While application of gabazine reduced the fractional difference in ON subfield sizes relative to control conditions, s-DSGCs * retained RFs that were 1.5-fold larger than the other ooDSGC types (Fig 5B, left).

**Figure 5.**
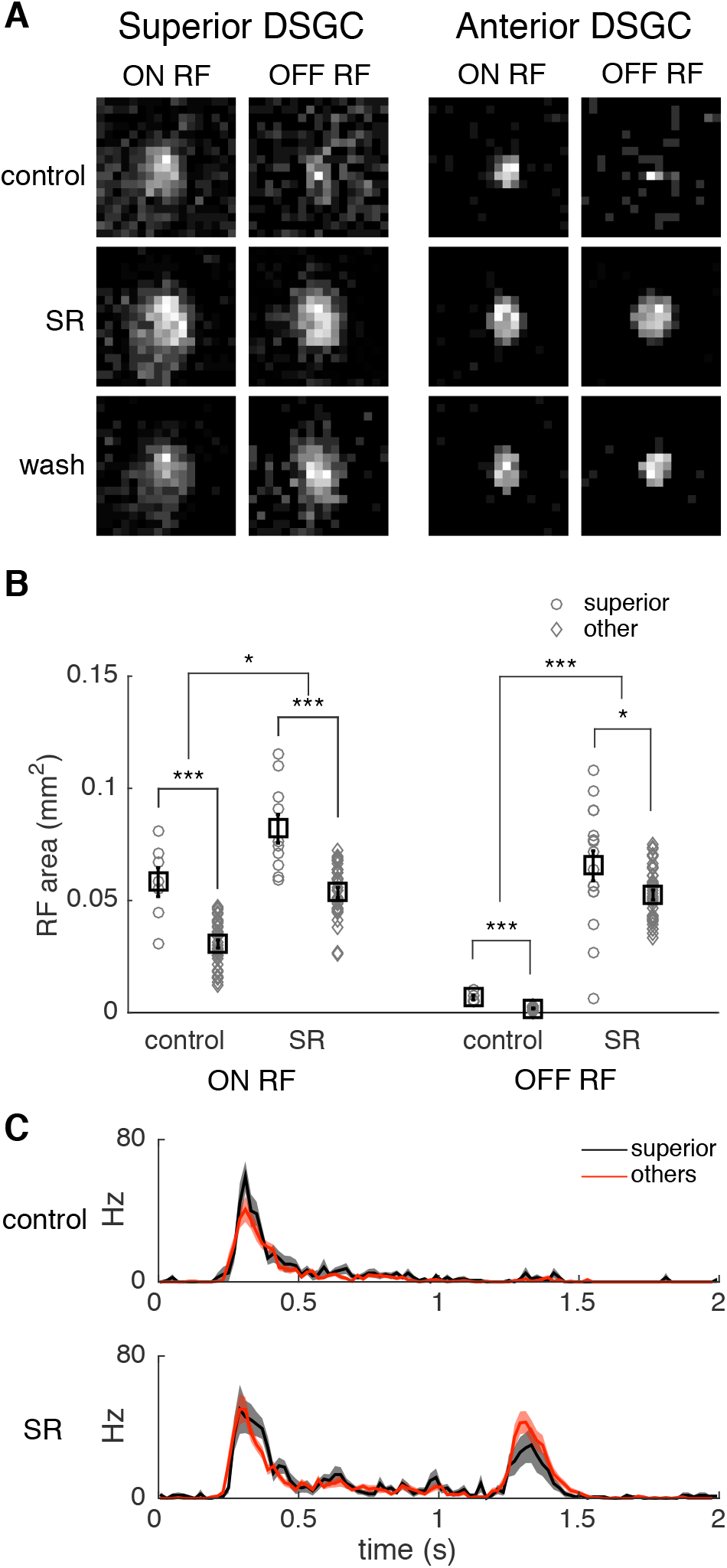
GABA_A_ blocker increases receptive field size of DSGCs. **A.** RF of a s-DSGC and other DSGC under control, SR and wash conditions. Stimulus square size: 120μm X 120μm, background light intensity: 0.2 R*/rod/s. **B.** Comparison of receptive field size across cell types and drug conditions for ON- and OFF responses, respectively (two retinas, superior: n = 17, others: n = 47). **C.** Responses of DSGCs to a single 120μm ×120μm square stimulus flashing at the center of their receptive fields. Center stimulus square was defined as the one that elicited the largest number of spikes for individual ganglion cells (one retina, superior: n = 12, others: n = 25, similar result observed in a 2^nd^ retina).

Previous work has shown that ooDSGCs are typically dominated by OFF responses at high light levels and prefer ON responses at lower light levels (Pearson and Kerschensteiner, 2015). Similarly, our results indicate that the OFF subfields become much smaller than the ON subfields under scotopic conditions (Fig 2, Fig 4B). Application of gabazine increased the OFF-subfield size (Fig 5B), and thereby unmasked a strong OFF response under scotopic conditions that was similar in amplitude to the ON response (Fig 5C). Thus, in addition to effective GABAergic inhibition controlling the absolute threshold and RF sizes of ooDSGCs, it controls response polarity by masking the OFF response under scotopic conditions.

## Discussion

We tested the hypothesis that s-DSGCs are more sensitive than other ooDSGC types near visual threshold. This hypothesis was based on previous work indicating that s-DSGCs exhibit broader tuning under scotopic conditions, and that this broader tuning may be advantageous for detecting motion across the population of ooDSGCs under dimly lit conditions (Yao et al., 2018). We show that the absolute sensitivity of s-DSGCs is ~10-fold greater than that of other ooDSGC types. This difference is largely mediated by differences in effective GABAergic inhibition with a smaller contribution from differences in RF area. We also show that GABAergic inhibition shapes the RF size and largely masks excitatory OFF input to ooDSGCs under scotopic conditions. Cumulatively, these results highlight a number of functional asymmetries across DSGCs, suggesting differences in the circuit mechanisms that produce their responses.

### What do these results suggest about DSGC circuits?

The circuit that provides input to DSGCs is perhaps the most intensely studied circuit in the retina. This is largely because the computation performed by the circuit is complex and precise, requiring finely-tuned developmental and synaptic mechanisms (Ding et al., 2016; Hanson et al., 2019; Huang et al., 2019; Poleg-Polsky and Diamond, 2016a, b; Sethuramanujam et al., 2016; Sethuramanujam et al., 2017). The circuit is also heavily studied because there are several mouse lines and molecular-genetic approaches to manipulate its development and function (Chen et al., 2016; Kay et al., 2011; Kim et al., 2010; Morrie and Feller, 2018). This technical leverage has produced a wealth of knowledge about the cell types that comprise the circuit. Nevertheless, our study provides data that are difficult to explain given previously identified circuit elements. This is probably because relatively few studies have examined DSGC physiology from starlight to sunlight and in a manner that distinguishes DSGCs based on their direction preference.

Our experimental approach produced two insights: First, s-DSGC (or their excitatory inputs) receive less GABAergic inhibition than other ooDSGCs (Fig 4). This observation predicts that there is a (yet to be identified) GABAergic amacrine cell that is engaged under low scotopic conditions that attenuates the spiking output of inferior, posterior, and anterior preferring DSGCs, but minimally attenuates s-DSGC responses (Fig 6A). Second, all ooDSGCs (or their OFF excitatory inputs) receive a source of inhibition that masks OFF responses under scotopic conditions (Fig 5). Again, this observation predicts another (yet to be identified) GABAergic amacrine cell with selective contacts onto either the outer dendrites of DSGCs and/or onto the OFF bipolar cells that provide input to these dendrites (Fig 6B). These putative amacrine cell types would need to be selectively engaged under scotopic conditions. We think it is unlikely that these inputs are coming from starburst amacrine cells because, (1) we have no evidence that the inhibition is direction tuned, (2) it needs to depend on light level, and (3) the inhibition must be differentially regulated between s-DSGCs and other ooDSGCs.

**Figure 6.**
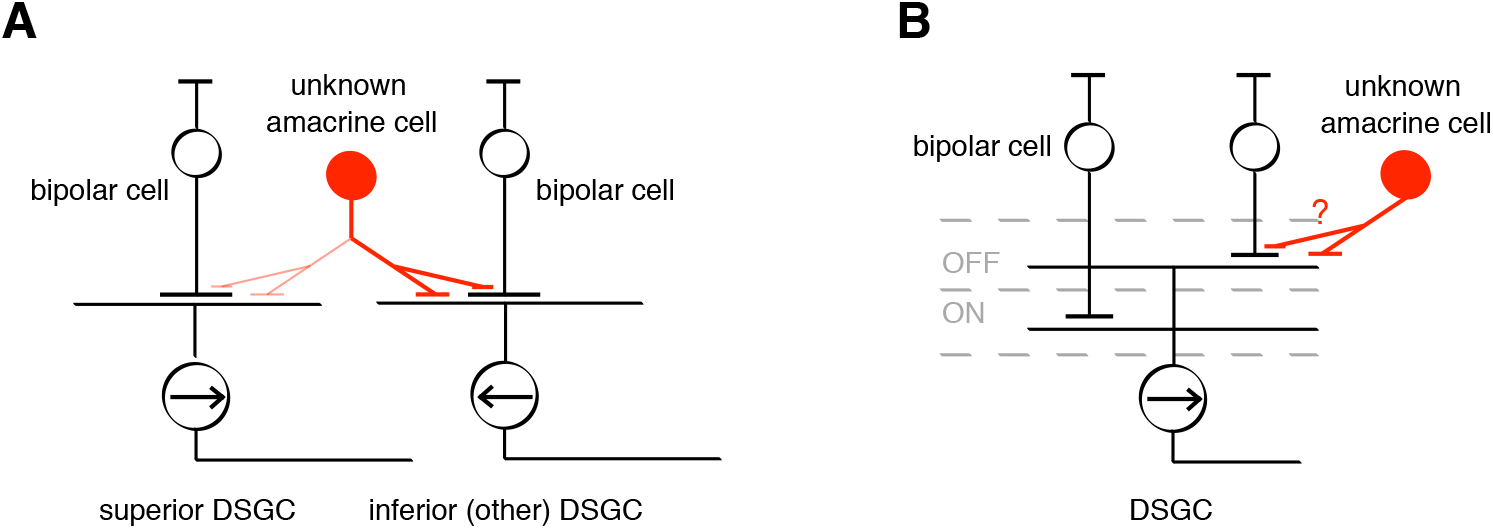
Synaptic organization between putative inhibitory amacrine cells and DSGCs. **A.** Schematic of possible circuity underlying differential adaptation across DSGCs. Under scotopic conditions, a putative amacrine cell type provides strong (thick red lines) GABAergic inhibition to the other ooDSGCs (or their presynaptic excitatory inputs) and weak (thin red lines) inhibition to s-DSGCs. **B.** Schematic of possible circuit underlying masked OFF responses of ooDSGC under scotopic conditions: A putative amacrine cell provides GABAergic inhibition to the OFF-layer dendrites of ooDSGCs and/or the OFF bipolar cells that provide excitatory input to ooDSGCs.

Finally, we emphasize these are hypothesized elements of the DSGC circuit that can account for our data. Proving their existence will require intracellular measurements that differentiate pre-versus post-synaptic inhibition and possibly a connectomics approach to identify novel cell types that are synaptically connected to DSGCs or their presynaptic partners (Bae et al., 2018; Ding et al., 2016; Helmstaedter et al., 2013; Kim et al., 2014).

### Inhibition limits RGC sensitivity

Different mouse RGC types exhibit distinct absolute sensitivities in darkness, with cells exhibiting threshold responses near one of four flash intensities: ~0.04, ~0.3, ~5, ~30 Rh*/rod (Pan et al., 2016; Volgyi et al., 2004). A recent study indicated that inhibition, putatively from amacrine cells, controls the absolute sensitivity of different RGC types (Pan et al., 2016). In particular, they observed that blocking GABA_C_ receptor-mediated inhibition with TPMPA lowered the absolute threshold of higher-threshold RGCs, making them similarly sensitive as the most sensitive RGCs. They also observed a minority of RGCs that exhibited lower thresholds when exposed to GABA_A_ receptor antagonist gabazine. It is difficult to know whether those RGCs sensitive to gabazine were DSGCs, because that study did not present moving stimuli (Pan et al., 2016). However, both Pan et al. 2016 and our study indicate that GABAergic inhibition masks high sensitivity responses in RGCs. Thus, it appears that many RGC types receive input through the high-sensitivity primary rod pathway, but that in some cases this input is masked by inhibition. The purpose of limiting these high-sensitivity signals to a minority of RGC types remains unclear. However, one possible explanation may come from our previous study on DSGCs, in which DSGCs that retained narrow and relatively high-precision DS tuning, were less sensitive to stimuli near threshold (Yao et al., 2018). It is possible that for RGCs that signal specific and complex visual features (e.g. motion direction) and are thus not acting as photon detectors, it is advantageous to only signal those features when the SNR of visual input is sufficiently high to accurately estimate the feature.

Blocking GABA_A_-mediated inhibition made the thresholds of s-DSGCs and other ooDSGC types more similar, but a clear threshold difference remained (Fig 4C). This residual difference with gabazine can be partly accounted for by the larger RFs of s-DSGCs (Fig 2), but according to our rod pool model, this only accounts for 30-50% of the residual threshold difference. The remainder is possibly the result of intrinsic differences in the signal-to-noise of the s-DSGCs or their synaptic inputs relative to the other ooDSGCs. Intracellular measurements under dark-adapted conditions are needed to further resolve these differences.

### Receptive field size and the influence of light adaptation

Under photopic conditions, we found that s-DSGCs exhibit RFs that are ~1.5-fold larger than other ooDSGC types (Fig 2F). This raises the possibility that either s-DSGCs have a higher coverage factor than other ooDSGCs types or they are lower in density. Density estimates of s-DSGC from the Hb9 line indicate ~80 cells/mm^2^ (Trenholm et al., 2011). Density estimates for posterior-preferring ooDSGCs from the DRD4 mouse line are 130 cells/mm^2^ (Rivlin-Etzion et al., 2011). This density ratio of 1.6 is similar to our observed RF size ratio of 1.5, suggesting similar coverage factors across ooDSGC types. Thus, DSGCs, like other RGC types appear to sample visual space with similar coverage under photopic conditions (Devries and Baylor, 1997; Gauthier et al., 2009a; Gauthier et al., 2009b).

The non-monotonic relationship between RF size and light level for DSGCs (Fig 2G) is likely partly shaped by GABAergic inhibition, which appears to reduce RF size below 1 Rh*/rod/s (Fig 5). We did not measure RF size across light levels under both control conditions and with gabazine, because such measurements would require more time and greater stability in the physiology than allowed for in our *ex vivo* experimental conditions. Thus, we have not fully mapped out the relationship between GABAergic inhibition, RF size, and light level. Another possible mechanism contributing to the non-monotonic relationship between RF size and light level is light-level dependent changes in homotypic coupling among photoreceptors, horizontal cells, and AII amacrine cells (Bloomfield and Volgyi, 2004; Jin and Ribelayga, 2016; Zhang et al., 2006). One interesting implication of these results is that the RF coverage factor across DSGCs is different between scotopic and photopic conditions. Assuming an RF coverage factor of one under photopic conditions, for s-DSGCs at 0.2 Rh*/rod/s, the increase in their RF size (Fig 2) indicates a coverage factor that approaches 3. However, for the other ooDSGC types at 0.2 Rh*/rod/s, their slight decreased in RF size suggests that coverage factor falls below one. Thus, while RF coverage factors are likely uniform (and ~1) across RGC types under photopic conditions (Devries and Baylor, 1997; Gauthier et al., 2009b), RF coverage factors appear to vary across RGC types under scotopic conditions.

### Role of inhibition in shaping parallel processing

Several recent studies indicate a crucial role for inhibition in differentially shaping parallel processing in the retina. First, inhibition in the outer retina from horizontal cells propagates to downstream circuit to have diverse influences on RGC function (Drinnenberg et al., 2018). Second, blocking inhibition in the inner and outer retina appears to largely eliminate differences in the output of 14 different bipolar cell types (Franke et al., 2017). Thus, the absence of inhibition reduces 14 distinct visual signals down to a uniform signal. Third, direction tuning and orientation tuning among RGCs is largely shaped by asymmetric GABAergic inhibition (Barlow and Levick, 1965; Hanson et al., 2019; Venkataramani and Taylor, 2010). This study complements these observations from the perspective of light adaptation. Specifically, adaptational differences across subtypes of DSGCs is largely caused by differences in effective GABAergic inhibition. Furthermore, GABAergic inhibition can differentially control the response polarity, RF size, and absolute sensitivity of RGCs at low light levels.

## Acknowledgements

We thank G. Awatramani for providing the FACx mice. We thank S. Roy, J Cafaro, and K. Ruda for commenting on drafts of this manuscript. The work was supported by NIH/NEI grant R01 EY024567 (G.D.F).

